# Comparative gene regulatory network mapping of Brassicaceae members with differential drought tolerance

**DOI:** 10.1101/2025.08.24.668636

**Authors:** Ramakrishnan Pandiarajan, Chung-Wen Lin, Mayra Sauer, Simin T Rothballer, Nora Marin-de la Rosa, Patrick Schwehn, Eleftheria Papadopoulou, Benedikt Mairhörmann, Pascal Falter-Braun

## Abstract

Climate change is projected to intensify weather extremes like droughts that threaten global crop yields. While many crops are sensitive to water deprivation, they have drought-tolerant wild relatives; however, the molecular basis for this differential tolerance among closely related species remains unclear. We previously investigated this using transcriptomics in the Brassicaceae family, comparing the drought-sensitive reference plant *Arabidopsis thaliana (Ath)* with its drought-tolerant relatives *Arabidopsis lyrata (Aly)* and *Eutrema salsugineum (Esa)*, uncovering key differences that may underlie their respective phenotypes. Here, to elucidate the regulatory architecture driving these differences, we mapped drought-responsive gene regulatory networks (GRNs) for all three species using the yeast one-hybrid (Y1H) approach. We cloned promoters of orthologs with divergent drought-induced expression patterns and screened them against the *Ath* transcription factor (TF) collection, generating a quality-controlled cross-species drought-responsive GRN map encompassing 1,137 high-confidence TF-promoter interactions. Comparative analysis revealed that the tolerant species share greater regulatory similarity and exhibit higher network connectivity than *Ath*. An *Esa*-specific expansion of bZIP TF interactions was observed, consistent with an enrichment of G-box motifs in its genome and in drought-upregulated genes. Several observed changes affect the ABA signalling pathway: *Ath* gained a susceptibility linked GBF3-*CYP707A1* edge, whereas *Esa* acquired tolerance associated ABF-*mTERF10* interactions. Furthermore, network-rewiring analysis uncovered a novel role for *ASIL2* in stress response. Finally, our network highlights TFs (CSDP1, ERF4, HB6, and MYB73) that likely contribute to the “stress-primed” state of *Aly* and *Esa*. Our work provides a broadly applicable framework for comparative GRN mapping and a valuable resource for improving drought tolerance in crops.

## Introduction

Climate change is a major threat to global food security. Shifts in freshwater availability and rainfall patterns are expected to cause more frequent and severe droughts, leading to declining crop yields and increased food insecurity. Since 2017, droughts have inflicted an estimated USD 124 billion in economic losses worldwide^1^. To mitigate these impacts and maintain crop productivity, drought-tolerant crops are required that can thrive under increasingly scarce and variable water conditions.

Drought tolerance relies on the precise coordination of abscisic acid (ABA)-dependent and - independent signalling, stomatal regulation, osmoprotectant synthesis, and the optimization of photosynthesis, growth, and metabolism. Most elite crop cultivars are highly drought sensitive, having been bred for maximum yields under ample irrigation^2^. By contrast, strong drought tolerance persists in wild relatives of major crops, such as *Solanum pennellii*^3^ (tomato) and *Oryza australiensis*^4^ (rice). This illustrates the existence of phenotypic plasticity of stress responses within plant families, i.e., tolerance traits appear and disappear within closely related family members.

To dissect the molecular basis of this plasticity, we chose the Brassicaceae as a model. The family includes the drought-sensitive reference plant *Arabidopsis thaliana (Ath),* and two morphologically similar but more drought-tolerant species, namely *Arabidopsis lyrata (Aly)* and *Eutrema salsugineum (Esa)* (Fig. 1a). Extensive genetic resources available for *Ath*, combined with the tractable laboratory growth of all three species, make this an ideal system for comparative analysis.

**Figure 1:**
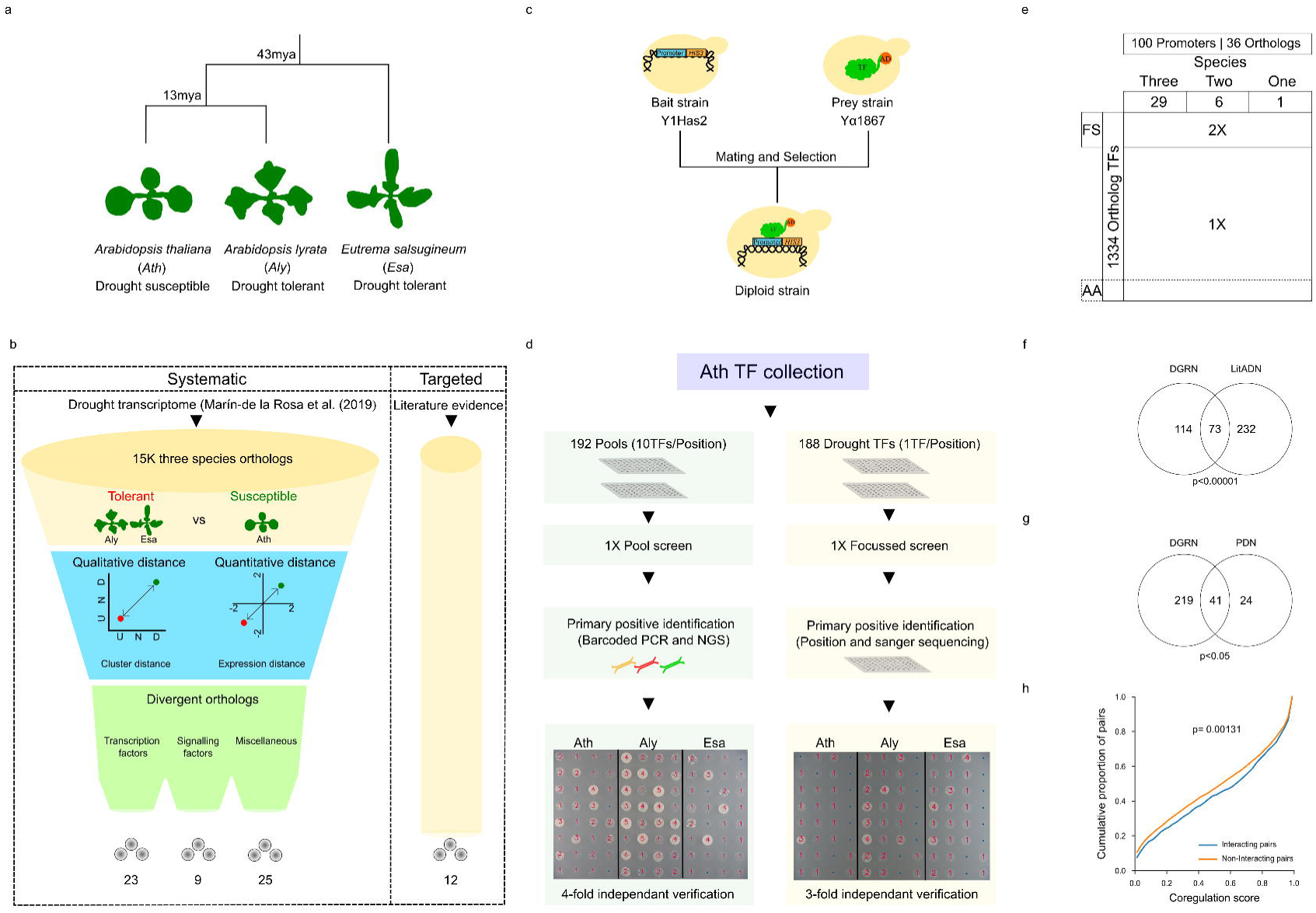
Generation of DGRN and its quality assessment. (a) Phylogenetic relationships among *Ath*, *Aly,* and *Esa.* (b) Schematic illustrating the orthologous promoter selection strategy used across the three species. (c) Diagram of the yeast-one-hybrid system (d) Overview of the pooled and focused Y1H screening pipelines. (e) Diagram of the DGRN search space, FS-Focussed screen, AA-Autoactivating TFs. (f) Overlap between the DGRN-*Ath* and *Ath* LitADN. (g) Overlap between the DGRN and PDN. (h) Distribution of coregulation scores for interacting TF-promoter pairs compared to non-interacting pairs.

In our earlier work, both *Aly* and *Esa* outperformed *Ath* under moderate drought^5^. We used developmentally synchronised *Ath*, *Aly* and *Esa* to collect gene expression data from rosettes at multiple time points, starting from the day watering was stopped (T0) and continuing up to 14 days later (T5, T11, T14), under both drought and well-watered conditions, and analysed 15,883 one-to-one orthologous genes shared across all three species. Time-series transcriptome revealed that *Aly* mounts an earlier and stronger response to drought, while *Esa* maintains a constitutively “primed” state with moderate transcriptional adjustments. Although both species achieve similar drought survival, their transcriptional programs differ markedly. The molecular mechanism underlying this distinct drought-coping strategy remains unclear. As transcriptional programs are ultimately controlled by gene-regulatory networks (GRNs), i.e. the transcription factors (TFs) binding to and regulating target promoters, we set out to map and compare the drought-responsive GRNs of *Ath*, *Aly,* and *Esa*, to reveal mechanisms underlying their differential drought tolerance.

Multiple strategies have been used to map stress-response GRNs. In a purely computational approach time series transcriptome data are converted into co-expression-based GRNs assuming a time-lag between TF and its downstream regulated genes. This method has revealed regulatory circuits involved in *Marchantia polymorpha* salt stress response^6^, *Ath* leaf-specific drought-stress response^7^, and the mannitol stress response^8^. However, co-expression GRNs rely on statistical correlations and are limited in power to establish direct regulatory causality. Experimental methods overcome these constraints. TF-centric assays such as ChIP-seq and DAP-seq identify promoters bound by a TF of interest, but they are less suited for promoter-centric questions. By contrast, the yeast-one-hybrid (Y1H) assay enables screening of promoters of interest against an entire TF collection, providing a systems-level view of its regulators. Y1H has been successfully used to unravel plant GRN circuits governing nitrogen metabolism^9^, phenolic metabolism^10^, defense regulation^11^, secondary cell wall synthesis^12^, regeneration^13^, vascular development^14^, ABA^15^, and auxin signalling^16^. With genome-wide ORFeome and TF collections now available for several plant species,^17–20^ comparative GRN mapping by Y1H is broadly applicable across plant species. Therefore, we chose a gene-centric Y1H approach to map physical TF-promoter interactions underlying drought responses in *Ath*, *Aly*, and *Esa*.

In this study, we build on our earlier drought transcriptome analysis by selecting genes with divergent expression and identifying their TF regulators through Y1H. Here, we present an experimental cross-species and gene-centric map of drought-responsive GRNs in the Brassicaceae. Benchmarking against independent TF-promoter interaction datasets confirms the high quality of our data. Comparative network analysis reveals topological similarities between the two tolerant species, Species-specific TF-promoter interaction biases, and key edge-rewiring events that shape species-specific expression patterns during drought. Collectively, these insights help understanding how closely related species evolve differential drought tolerance.

## Results

### Mapping of drought responsive GRN

To investigate the molecular basis of differential drought tolerance between *Ath*, *Aly,* and *Esa*, we scored 15,883 orthologous genes in all three species for divergent expression patterns during drought^5^. Divergence scores combined both qualitative (up vs down-regulation) and quantitative (fold-change) information (Fig. 1b). We focused on earlier expression differences at T5 and T11 and excluded T14, as *Ath* showed a marked loss of viability upon re-watering after day 14. For T5 and T11, each three-way ortholog was assigned a qualitative descriptor: upregulated (U), downregulated (D), or non-regulated (N) relative to the preceding time point, yielding nine possible regulatory patterns (UU, NN, DD, UN, UD, NU, ND, DU, DN) (Fig. 1b). To capture tolerant species-specific patterns, we only scored *Aly*-specific, *Esa*-specific, and *Aly*-*Esa* common expression divergence relative to *Ath*. We scored and grouped regulatory proteins, such as TFs and kinases, separately (Fig. 1b), as their expression changes may be amplified through downstream signalling and therefore even relatively modest differences may have a notable phenotypic impact. Overall, this systematic procedure yielded a core set of 110 orthologs with the most divergent drought-responsive expression patterns. We supplemented this data-driven set with two drought stress markers (RD29B and LTP4) and 18 genes involved in ABA signalling and proline metabolism, thereby bringing the total to 130 three-way ortholog sets. We then set out to map the GRN driving the divergent expression pattern of these orthologs by Y1H (Table S1).

To clone promoters of the 130 divergent orthologs, we surveyed publicly available *Ath* TF-binding motifs across the -3000 to +1000 bp window relative to the start codon (ATG). As TF motif density peaked between -1300 and +300 bp, we designed primers to amplify fragments of this length for each ortholog. When suitable priming sites were absent, short primer-walking was done upstream and downstream of ATG. We ensured that the promoters from all three species within an ortholog set covered the same region and were of the same size. PCR products were cloned into the pMW2 vector upstream of the auxotrophic reporter *HIS3*, which confers the yeast strain, ability to grow without histidine relative to its expression levels (Fig. 1c). The pMW2 vector was linearised and integrated into the genome of haploid yeast, generating promoter bait strains. This workflow yielded bait strains for 100 promoters: 87 representing complete sets of 29 three-way orthologs and 13 covering one or two species per ortholog set (Fig. 1e). Collectively, these strains harbouring the promoters of all three species enabled us to proceed towards a systematic Y1H screen to identify their upstream TF regulators.

A comprehensive Y1H screen requires a complete set of TFs from the organism under study. Such a resource is only available for *Ath* ^21^, but not for *Aly* and *Esa*. We therefore used the *Ath* TF collection to screen promoters from all three species. This approach is justified by extensive evidence showing that TF-DNA binding specificities are broadly conserved across large evolutionary distances. A survey spanning 54 different DNA binding domain (DBD) classes from 131 eukaryotes showed that closely related DBDs almost always recognise similar DNA motifs^22^. In plants, a comparative DAP-seq analysis of four Brassicaceae species, including *Ath* and *Esa*, revealed highly congruent binding preferences for ABA-related TFs^23^. Crop TFs also function interchangeably when overexpressed in *Ath* and other species^24,25^. Given this conservation, the TF-prey library for cross-species TF-promoter interaction mapping was generated by introducing *Ath* TF-GAL4 activation domain fusions into haploid yeast.

TF-promoter interactions were screened by mating haploid promoter-bait strains with haploid TF-prey strains and subsequent scoring of growth based on *HIS3* selection (Fig. 1d). To screen efficiently, we opted for a dual strategy: an unbiased pool screen and a focused 1-to-1 screen (Fig. 1d). While the pool screen covered the entire collection, the focused screen restricted to TFs annotated with drought-related GO terms increased the saturation for the most relevant regulators. Unexpectedly, we observed a high prevalence of TFs that bound the pMW2 plasmid backbone carrying the promoter insert, a phenomenon not previously reported. To remove these artefacts (Fig. S1a), we compared TF binding to two backbone controls integrated in yeast: a complete pMW2 vector containing the Gateway ccdB cassette and an empty pMW2 vector lacking the cassette. Interactions from TFs that bound to both controls (n = 96) were discarded to eliminate false positives. To minimise false negatives, due to incomplete saturation of the screen, any interaction detected for a promoter in one species was retested with orthologous promoters from the other two species. Candidate TF-promoter pairs were tested alongside empty-vector and GFP negative controls, and yeast growth was quantified with a convolutional neural network model^26^. Pairs were considered positive when yeast growth exceeded the negative control by ≥ 2 units on a 0-5 scale. We only considered pairs as bona fide interactions if they were reproduced in at least 2/3 (focused screen) or 3/4 replicates (pool screen), respectively. After this stringent filtering, we retained interactions involving 1,334 TF with orthologs in all three species, yielding 1,137 TF-promoter pairs that define the drought-responsive GRN (hereafter DGRN) for the species under study.

### Validation of DGRN using orthogonal assays

To assess the quality of our data^27^, we compared DGRN with both literature and experimental datasets. First, we benchmarked our DGRN-*Ath* network against the literature-derived *Ath* DAP-seq network^28^ (LitADN). Among the TFs common to both datasets, DGRN-*Ath* contained 187 interactions, while LitADN had 305. A total of 73 interactions were shared between the two networks, representing a statistically significant overlap (P < 1 × 10^-4^) when compared to 10,000 degree-preserving random rewired versions of DGRN-*Ath* (Fig. 1f).

Next, we tested a subset of our interactions experimentally with a modified ampDAP-seq^29^ protocol. Instead of amplified genomic DNA, we used PCR-amplified promoter fragments from our clones as a DNA library and assayed them against 14 *Ath* proteins expressed in *E. coli*. High-throughput sequencing of TF-bound amplicons yielded binding peaks that delineated both the location and intensity of TF-promoter interactions. Motifs derived from these peaks matched published motifs for several TFs (Fig. S1b). Peak distribution across each promoter was also informative: the transcriptional repressor ethylene response factor 4 (ERF4) bound predominantly within the coding region, whereas the activator ABA binding factor 4 (ABF4) bound in the upstream promoter region (Fig. S1b). Thus, promoter-ampDAP reliably captures TF-DNA binding. For the five proteins that produced high-quality peaks (Fc ≥ 5), we obtained 65 TF-promoter interactions, forming the promoter-ampDAP network (PDN). Comparison of PDN with our Y1H-derived DGRN revealed 41 shared interactions, a significant overlap (P = 3.67 × 10^-2^, Fisher’s exact test, Fig. 1g). Importantly, this 15% validation rate falls within the 11-20% range reported for previous high-quality Y1H studies (Table S1).

We next assessed the functional relevance of the DGRN by checking if TF binding to a promoter translates into changes in the target gene’s expression. We devised a coregulation score (Methods) that calculates the coefficient of determination between log_2_FC values of each TF and its target promoter across three-time drought points, capturing both positive and negative regulation in a single metric (Fig. 1g). Interacting TF-promoter pairs in the DGRN had significantly higher coregulation scores than the non-interacting pairs (P = 1.3 × 10^-3^, Mann-Whitney U Test). Thus, DGRN interactions are associated with regulatory gene expression changes, supporting the functional validity of the network.

Finally, we aimed to experimentally demonstrate whether identified TF-promoter interactions lead to regulatory gene expression changes. We employed a tobacco transactivation assay^30,31^ with minor modifications using the cloned full-length promoters and the TFs. Notably, the TFs, Arabidopsis transcriptional activation factor 1 (ATAF1) and ERF4, activate and repress the promoter they bound in our transactivation assay, demonstrating its utility. We tested if binding leads to a regulatory outcome in eight random interactions from the dataset involving pairs from all three species. Five of the eight tested interactions had a significant regulatory outcome (Fig. S1c), confirming not only that the dataset is of high biophysical quality but also that TF-promoter pairs in DGRN are functionally meaningful *in planta*.

Collectively, these analyses indicate that DGRN interactions considerably overlap with literature and experimentally determined interactions, highlighting the regulation and specificities of TF-promoter binding, reinforcing the high quality of our data, and prompting further thorough analysis.

### Characteristics of DGRN

The complete DGRN map encompasses 374 *Ath* interactions (35 promoters), 366 *Aly* interactions (31 Promoters), 397 *Esa* interactions (33 promoters) (Fig. 2a). For 29 complete three-way ortholog sets, 996 interactions with 81 TFs were identified. Among these sets, all promoters, except the *Esa* NAC41-promoter, yielded at least one interaction demonstrating the functionality in the Y1H assay. Several promoters and TFs in the DGRN were linked to ABA signaling and diverse abiotic/stress responses. Many uncharacterized genes were also identified, highlighting the potential of our approach to uncover novel stress regulatory roles (Fig. 2b).

**Figure 2:**
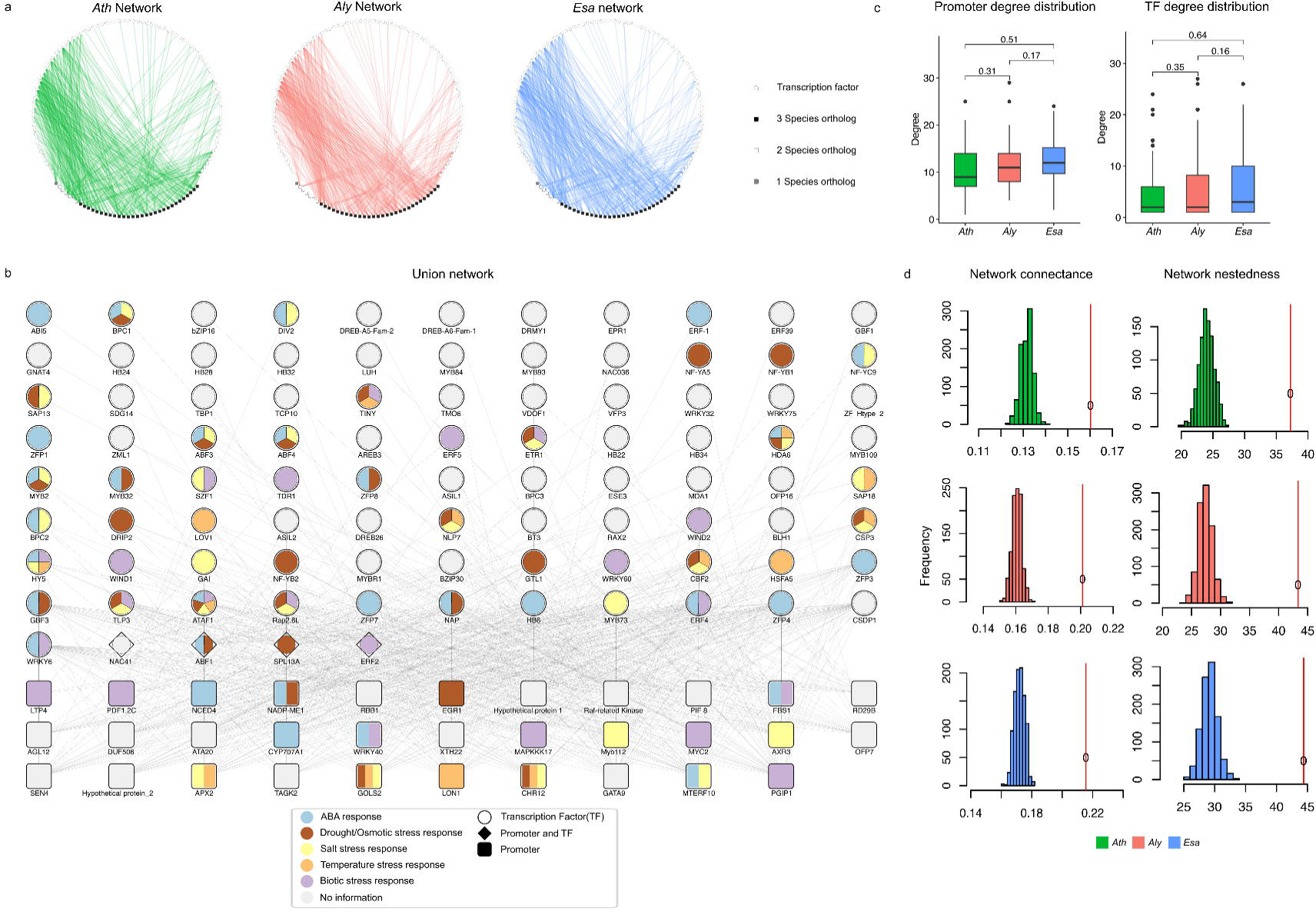
DGRN overview. (a) Complete DGRN comprising *Ath*, *Aly,* and *Esa* networks. (b) Union network showing the combined set of unique TF-promoter interactions across all three species. (c) Distribution of promoter and TF degree in all three species. (d) Comparison of connectance and nestedness of all three species networks.

In our dataset, each promoter averaged 11.5 incoming TF edges (median = 11), and each TF had an average outdegree of 12.2 promoters (median = 3) (Fig. 2c). Despite differences in the assay setup and promoter length utilized in our assay, these observed in- and out-degrees are similar to previously published Y1H networks^32^. Since we used *Ath* TFs for *Aly* and *Esa* promoters in our screen, we checked if there were global differences in the promoter/TF degree distribution patterns across the species (Fig. 2c). Overall, no species-specific differences were observed, suggesting that *Ath* TFs indeed function well with *Aly* and *Esa* promoters in our system. Consistent with the conserved binding specificities of closely related TFs^22^, members of the same family frequently targeted overlapping promoter sets; notable examples include the Basic Penta Cysteine (BPCs), trihelix Arabidopsis 6B Interacting Protein 1-Like (ASILs), and ABF families (Fig. S1d). These findings suggest that plant TFs have retained their binding preferences in our assay and do not discriminate among orthologous promoters from *Ath*, *Aly*, and *Esa* when the appropriate binding sites are present.

We examined the overall network topology in each species using the complete 29-ortholog set network to investigate tolerant and susceptible species networks. Two network parameters stood out: connectance (the fraction of all possible TF-promoter edges that are observed) and nestedness (the extent to which connections are hierarchically structured). Networks with higher connectance and nestedness exhibit greater resistance to simulated interaction losses^33^. We found that both drought-tolerant species had higher connectance (*Aly*-0.20, *Esa*-0.21) and higher nestedness (*Aly*-43.3, *Esa-*44.4) than the susceptible *Ath* (connectance = 0.16, nestedness = 37.2) (Fig. 2d), relative to randomized networks. These findings suggest that the drought-regulatory circuits in tolerant species are intrinsically more connected and nested. Whether this is a global feature of their stress response or a consequence of our limited promoter selection needs to be clarified by future studies.

### Network divergence trends in DGRN

We wondered if the overall interaction patterns of the three species reflect their (evolutionary) sequence similarity (*Ath-Aly* closest) or drought response (phenotypic) similarity (*Aly-Esa* closest). Across the 29 complete ortholog promoter sets, the *Ath-Aly* pair expectedly showed the highest average sequence similarity (68%) in comparison with the more distantly related species pairs; *Aly-Esa* 57% ( P = 5.12 × 10^-5^, Mann-Whitney U-test) and *Ath-Esa* 61% ( P = 1.48 × 10^-4^, Mann-Whitney U-test). We then compared this observation with interaction similarities by calculating the Jaccard Index (JI) for all species pairs across the 29 complete ortholog sets (Fig. 3a). Remarkably, in this analysis, the *Aly-Esa* pair, involving the two drought-tolerant species, had the highest similarity (Mean JI = 0.47) compared to *Ath-Esa* (Mean JI = 0.37, P = 0.02, Student’s t-test) and *Ath-Aly* (Mean JI = 0.38, ns) (Fig. 1a). These overall patterns are exemplified by *Hypothetical protein 1*, for which TF-promoter interaction similarities (Fig. 3b) are also mirrored in transcriptional responses to drought. While its transcript is upregulated early in both *Aly* and *Esa* (T5) following the onset of drought stress, the regulation in *Ath* occurs only in the final stage of drought stress at T14 (Fig. 3c). Thus, rather than tracking evolutionary distance, TF-promoter interaction patterns align more closely with shared drought-tolerance phenotypes.

**Figure 3:**
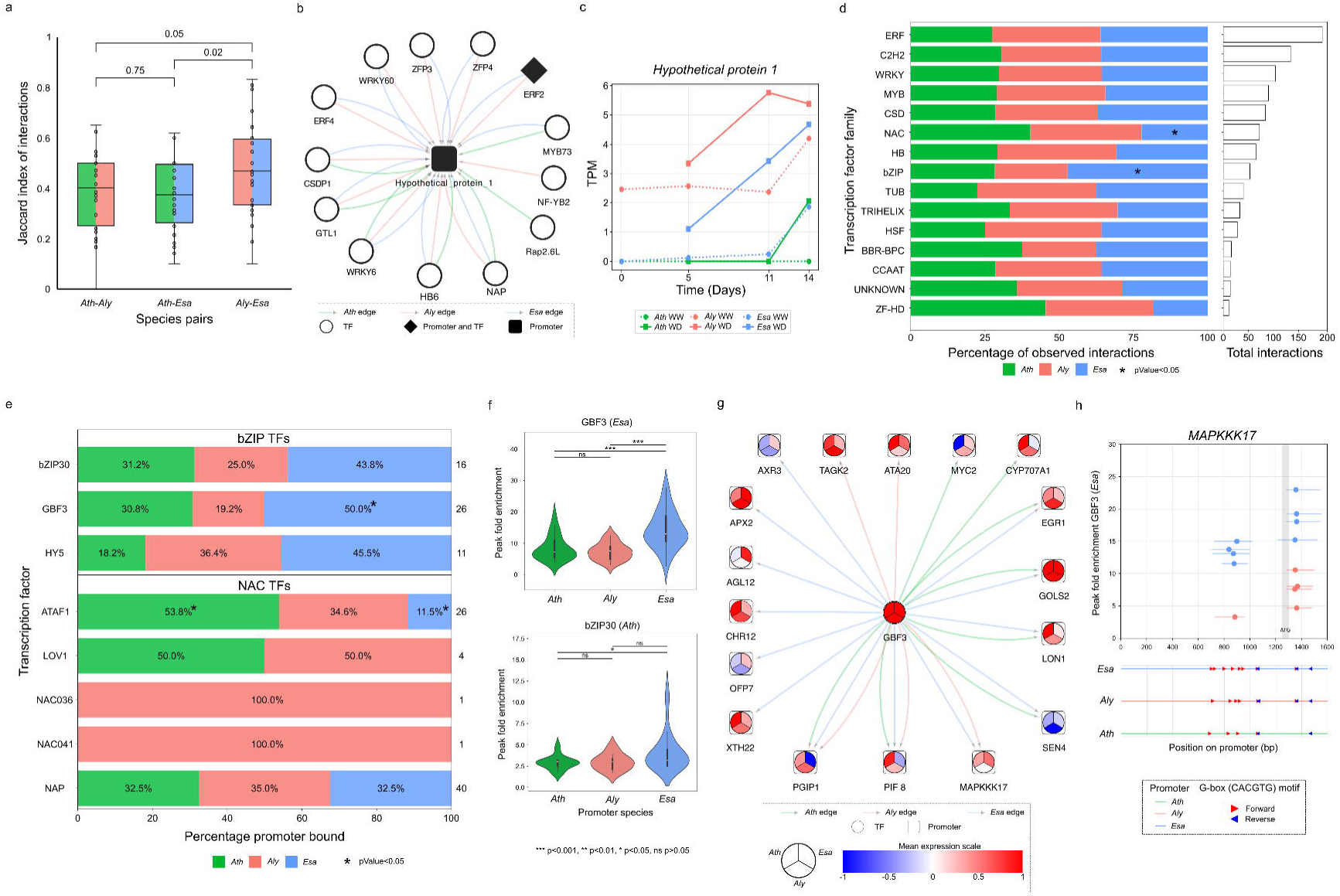
DGRN rewiring trends. (a) Jaccard similarity index comparing TF-promoter interaction overlap between species pairs. (b) *Hypothetical protein 1* regulatory network. (c) Expression profile (TPM) of *Hypothetical protein 1* during the drought time-course experiment. (d) Interaction patterns of TF families within DGRN (families with ≥10 interactions shown). (e) Interaction profiles of NAC and bZIP TFs across the DGRN. (f) Distribution of pampDAP peak fold enrichment distribution for GBF3 (*Esa*) and bZIP30 (*Ath*). (g) Regulatory network of GBF3. (h) pampDAP peak fold enrichment of GBF3 (*Esa*) across the promoter length of *MAPKKK17,* with G-box motif positions in all three species.

We next examined interaction trends at the TF level, starting with a family-wide survey. Expansion or contraction of TF families and the parallel turnover of their binding sites in the genome have been recognised as a key driver of phenotypic innovation in plants^34^. We therefore asked whether species-specific interaction biases exist among the 28 TF families present in our network (Fig. 3d) (raw P-values, no FDR correction). Indeed, members of the NAM, ATAF1/2, and CUC2 (NAC) family, which have been associated with senescence^35^, have increased interactions in *Ath* (P = 0.04, OR = 1.61, Fisher’s exact test), but decreased interactions in *Esa* (Only 16/72 interactions, OR = 0.52). In contrast, basic leucine-zipper (bZIP) factors, key mediators of ABA signalling and stress tolerance^36^, have increased interactions in *Esa* (P = 0.03, OR = 1.75, Fisher’s exact test). These observations suggest species-specific rewiring of stress-response regulation, aligning with the known roles of NACs and bZIPs in drought tolerance.

We next sought to pinpoint the individual NAC and bZIP TFs that drive the family-level biases (Fig. 3e). ATAF1, a NAC-family negative regulator of ABA signalling^37^ illustrates the trend: it had significantly fewer interactions in *Esa* (P = 5 × 10^-3^, OR = 0.17) but significantly more in *Ath* (P = 0.01, OR = 3.51, Fisher’s exact test). The pattern for bZIPs was even more pronounced: all three TFs present in the network, bZIP30, G-box binding factor 3 (GBF3), and elongated hypocotyl 5 (HY5), showed elevated interaction counts in *Esa*. The strongest signal came from GBF3, a positive regulator of ABA signalling and drought tolerance^38^, which had significantly more interactions in *Esa* (P = 0.04, OR = 2.77, Fisher’s exact test). To further support these findings, we performed a promoter-ampDAP (pampDAP) assay with *Esa* GBF3 and *Ath* GBF3; however, only *Esa* yielded results. As the *Esa* and *Ath* proteins are 100% identical in the DNA binding bZIP domain, and the extracted binding specificity of *Esa* GBF3 is identical to the published binding motif of *Ath* GBF3 (Fig. S1b), we consider the *Esa* GBF3 data not only informative but complementary to the *Ath* GBF3-based observations in Y1H. In pampDAP dataset, GBF3 binding peaks on *Esa* promoters were significantly stronger than other species (*Ath* vs *Esa*, P = 9.83 × 10^-5^; *Aly* vs *Esa*, P = 8.3 × 10^-6^, Mann-Whitney U test) (Fig. 3f). The same trend was observed for *Ath* bZIP30 (*Ath* vs *Esa*, P = 0.03, Mann-Whitney U Test). Thus, multiple lines of experimental evidence confirm a species-specific binding bias of bZIP family factors toward *Esa* promoters.

Next, we investigated how the *Esa*-bZIP binding bias manifests at the promoter level, focusing on GBF3-bound promoters (Fig. 3g). GBF3 recognizes the G-box core motif 5’-*CACGTG*-3’. A striking example is the orthologous promoters of *mitogen-activated protein (MAP) kinase kinase kinase 17 (MAPKKK17)*. The *Esa* promoter contains 11 G-box core motifs, compared to nine in *Aly* and six in *Ath* (Fig. 3h). Although the *Ath* promoter harbors binding motifs, GBF3 binding was detected only in *Esa* and *Aly,* in both DGRN and pampDAP-implying that motif number and spatial arrangement are critical determinants of binding. Detailed mapping revealed two clusters of closely spaced G-boxes: one proximal to the ATG and another more distal. *Ath* lacks one motif in the proximal cluster, whereas *Esa* retains both clusters, resulting in overall higher peak fold-enrichment in the pampDAP. Recent work indicates that arrays of low-affinity sites can merge into high-affinity energetic sinks, enhancing TF recruitment and transcriptional output^39^. Our data is consistent with this model, as high fold-enrichment peaks of GBF3 align precisely with the G-box clusters in *Esa*. Functionally, MAPKKK17 is a key activator of the MAPK cascade in ABA signaling. It is induced by ABA and osmotic stress in an ABA-core-dependent manner, acts upstream of MAP kinase kinase 3 (MKK3), and ultimately activates MAP kinase 7 (MPK7). Both loss-of-function *mkk3-1* and *mapkkk17mapkkk18* double mutants exhibit increased water loss, underscoring the module’s importance for drought tolerance. Thus, we propose that a species-specific expansion of bZIP TF binding sites in the *Esa MAPKKK17* promoter enhances ABA sensitivity and constitutes network rewiring that contributes to the improved drought tolerance of *Esa*.

We asked whether the expansion of GBF3 binding sites observed in *MAPKKK17* promoter reflects a genome-wide trend that enhances the drought response in *Esa*. A whole-genome motif scan revealed that a significantly larger proportion of *Esa* promoters contain the G-box core motif (18.52%) than *Ath* promoters (17.42%) (P = 9.84 × 10□□, Chi-square test). To determine whether this difference translates into transcriptional output, we compared G-box core motif frequencies in genes that are upregulated (log_2_FC ≥1) and downregulated (log_2_FC ≤-1) during drought stress^5^. Indeed, motifs of the transcriptional activator GBF3 were enriched in upregulated genes (P = 1.09 × 10□^4^, Fisher’s exact test) and depleted in downregulated genes (P = 2.89 × 10□^3^, Fisher’s exact test) in *Esa* relative to *Ath*. Thus, the expansion of GBF3 binding sites appears to be an important aspect of *Esa*’s enhanced transcriptional response to drought, contributing to its improved drought tolerance.

### Differential drought signaling in DGRN

To pinpoint regulatory differences that might underlie the species-specific drought responses, we examined the ABA signaling pathway. ABA is the central hormonal signal by which plants sense and respond to drought. It is synthesized by 9-cis-Epoxycarotenoid dioxygenase (NCED) enzymes and perceived, in various cell types, by the RCAR/PYR/PYL family of receptors. These receptors act as regulatory subunits of group A type-2C protein phosphatases (PP2Cs), which activate SNF1-related protein kinases 2 (SnRK2s). Activated SnRK2s phosphorylate AREB and ABF TFs, thereby driving ABA-dependent transcriptional response^40^. ABA levels are counter-balanced by catabolic enzymes, such as members of the CYP707A family^40^.

To identify DGRN edges that could mediate differential drought tolerance via altered ABA signaling, we systematically annotated every node as either a positive or negative regulator of drought stress tolerance using published data on ABA responses and desiccation stress phenotypes (drought, osmotic, and salt stress). A node was labelled as positive if (i) its overexpression increased ABA sensitivity or its loss of function reduced ABA sensitivity, and/or (ii) its overexpression enhanced stress tolerance or its loss of function increased stress susceptibility. Conversely, a node was labelled as negative if (i) its overexpression reduced ABA sensitivity or its loss of function increased ABA sensitivity, and/or (ii) its overexpression increased stress susceptibility, and its loss of function improved stress tolerance. Utilizing these annotations, we were able to identify several edge rewiring events that can be related to the differential tolerance phenotypes of the species under study.

Intriguingly, we identified an *Ath*-specific link between the transcriptional activator and positive stress response regulator GBF3 to the promoter of *CYP707A1*, an ABA-degrading enzyme that acts as a negative tolerance regulator. This edge coincides with the presence of an additional GBF3 binding site in the *Ath CYP707A1* promoter when compared to *Esa*. Consistent with this *Ath* specific edge, *CYP707A1* is upregulated throughout the drought period only in *Ath*, mirroring the expression pattern of GBF3 (Fig. 4a). Although GBF3 is induced in both *Esa* and *Ath,* it is connected to an ABA-inactivating gene solely in *Ath* revealing species-specific rewiring of ABA signalling that may contribute to *Ath* drought susceptibility.

**Figure 4:**
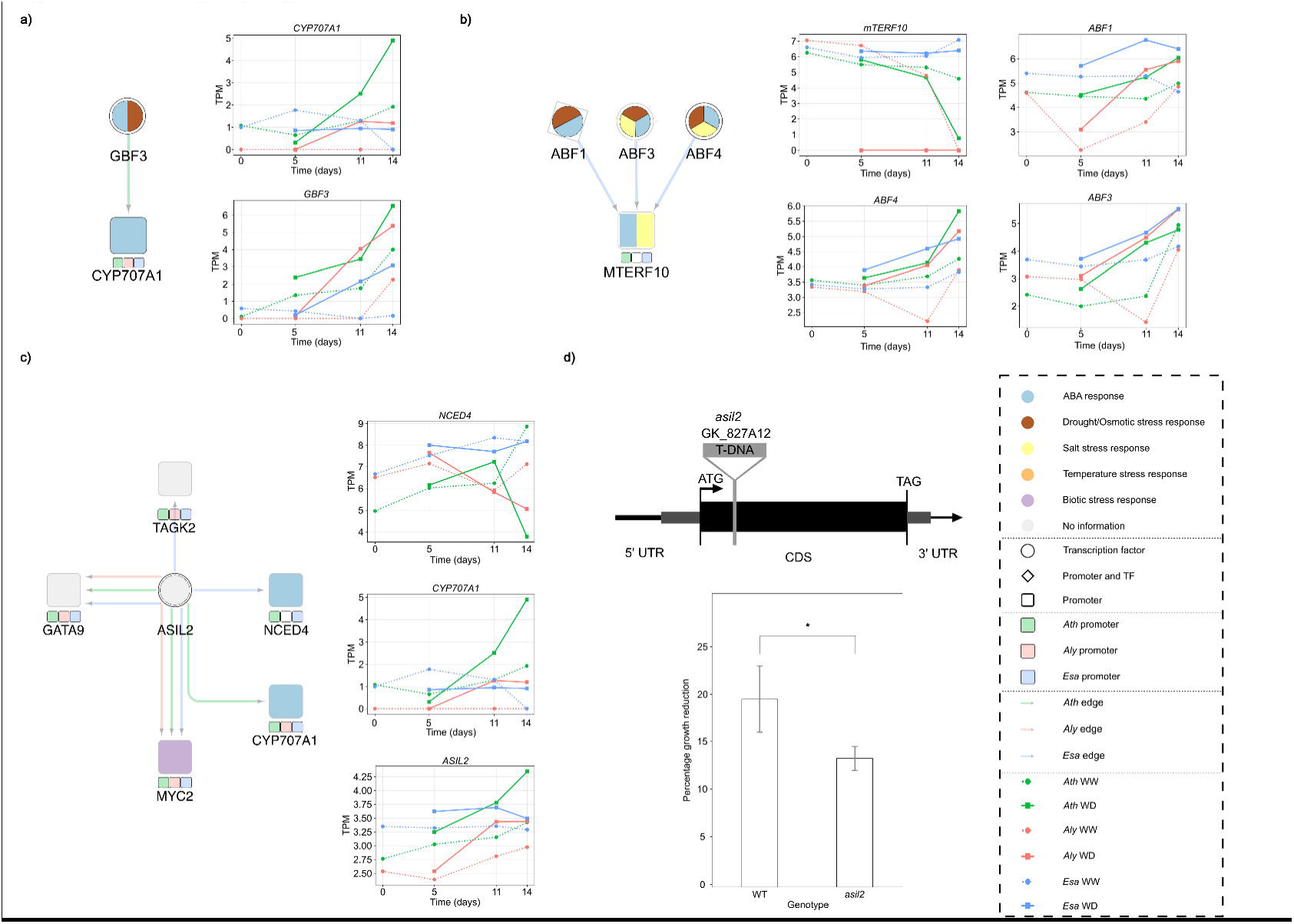
Differential drought signalling in DGRN. (a) Rewiring of the GBF3-*CYP707A1* edge and corresponding gene expression (TPM) under drought. (b) Rewiring of the ABF-*mTERF10* edge and expression (TPM) under drought. (c) ASIL2 regulatory subnetwork and expression (TPM) dynamics under drought. (d) Phenotypic response of the *asil2* mutant on 25 mM Mannitol; * p< 0.05

A second example involves *mitochondrial transcription termination factor 10 (mTERF10),* a positive regulator of salt stress tolerance^41^. In drought-tolerant *Esa* but not in susceptible *Ath*: the ABA responsive TFs ABF1, ABF3, and ABF4 bind the *mTERF10* promoter (Fig. 4b). This interaction pattern matches the gene’s expression profile during drought: *mTERF10* transcript levels remain stable in *Esa* but decline progressively in *Ath* (Fig. 4b). ABFs likely maintain the sustained expression levels of *mTERF10* in *Esa*. As these ABFs are transcriptional activators expressed at elevated levels throughout drought in *Esa* relative to *Ath*, they may counteract any repressors acting on *mTERF10* promoter. Accumulating such new edges between two classes of positive regulators of tolerance-ABFs and *mTERF10*-might collectively strengthen stress signalling, contributing to enhanced stress tolerance in *Esa*.

A third example involves the trihelix transcription factor ASIL2, which represses embryo-maturation during early seed development^42^, but has not yet been linked to abiotic stress. Strikingly, in the drought-tolerant *Esa*, ASIL2 binds the promoter of the positive tolerance regulator and ABA-biosynthesis gene *NCED4*, whereas in *Ath* it targets the ABA-catabolic gene *CYP707A1*, a negative tolerance regulator (Figure 4c). In *Ath*, *ASIL2* transcripts accumulate throughout the drought, mirroring *CYP707A1* expression and suggesting a role in accelerating ABA degradation. Thus, it appears that ASIL2 may act to promote drought tolerance in *Esa* but counteract it in *Ath*. To test the latter, we analysed the *Ath* T-DNA line GK_827A12, which carries an exon insertion in *ASIL2* (Figure 4d). Seven-day-old seedlings were grown ± 25 mM mannitol to impose osmotic stress. After seven days of stress, wild-type plants exhibited a 20% reduction in rosette area, in comparison to 13% in the *asil2* mutant (P = 0.04, Student’s t-test). Thus, indeed in *Ath* ASIL2 functions as a negative regulator of osmotic-stress tolerance, likely by promoting the expression of *CYP707A1*.

### Temporal drought response changes in DGRN

Constitutive expression of stress-response genes can enhance plant tolerance^38^. In our earlier transcriptomic study, we observed that the tolerant species *Aly* and *Esa* maintain a “stress-ready” state even under well-watered conditions^5^; several tolerance genes already show higher basal expression before water deficit is imposed (T0). To identify TFs that contribute to this “stress-ready” state, we identified orthologous promoters whose basal expression at T0 was higher in the tolerant species than in *Ath* (log_2_FC ≥ 1, relative to *Ath*) and extracted their DGRN interactions. We then retained only those interactions involving TFs that bound at least five of the high basal expression promoters in *Aly* or *Esa*. This filtering produced two species-specific, T0-focused subnetworks: (i) an *Aly* subnetwork comprising seven TFs and seven promoters, and (ii) an *Esa* subnetwork comprising ten TFs and ten promoters (Fig. 5a).

**Figure 5:**
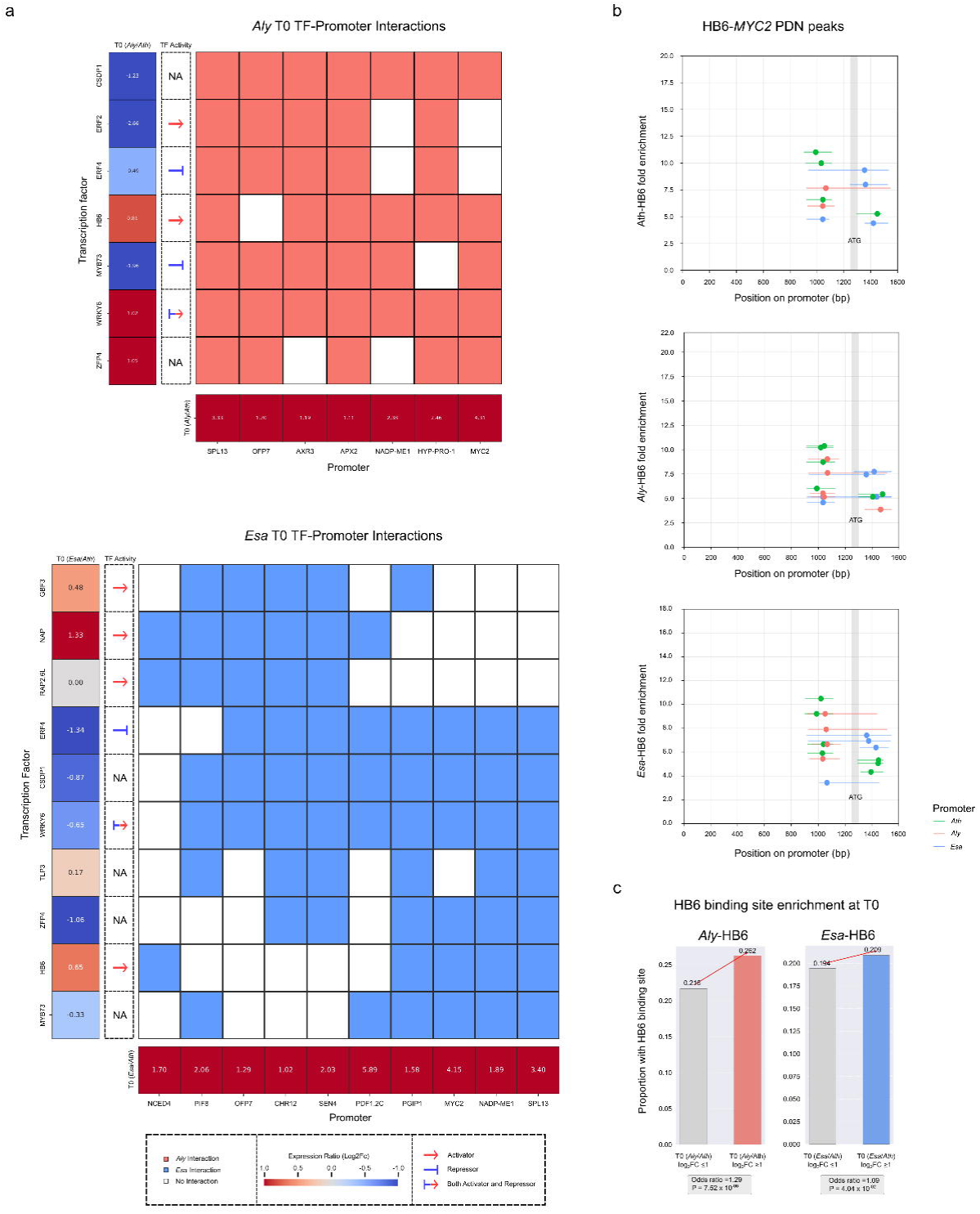
T0 *Aly* and *Esa* temporal network. (a) *Aly* and *Esa* T0 TF-promoter interaction subnetwork (Promoter orthologs with log_2_FC ≥ 1 relative to *Ath* shown). (b) pampDAP fold enrichment profile of HB6 (*Ath*, *Aly,* and *Esa*) on *MYC2* promoter. (c) Enrichment of HB6 binding sites in genes that are constitutively expressed in *Aly* and *Esa* relative to *Ath*.

Comparison of the two T0-focused subnetworks revealed four promoter orthologs shared by *Aly* and *Esa*: *MYC2, NADP-dependent malic enzyme 1 (NADP-ME1), squamosa promoter binding protein-like 13A (SPL13A),* and *ovate family protein 7 (OFP7).* All four orthologs remain expressed at elevated levels in both tolerant species relative to *Ath* throughout the drought period, pointing to a potential contribution to the ‘drought-ready state’. Functional evidence supports this interpretation. Overexpression of *MYC2* improves water-use efficiency in poplar^43^ and tomato^44^. SPL13 promotes cuticular wax biosynthesis^45^, and *Esa* leaves indeed contain more wax than those of *Ath*^46^, limiting non-stomatal water loss^47^. Cytosolic NADP-ME1 supplies NADPH for ROS scavenging; its rice ortholog confers broad abiotic-stress tolerance when expressed in *Ath*^48^, whereas *nadp-me1* mutants display attenuated ABA responses^49^. Although OFP7 is best known for regulating leaf and cotyledon shape^50^, its elevated expression in *Aly* and *Esa* may still influence water-use efficiency by altering leaf morphology, a hypothesis that remains to be tested. Collectively, the elevated expression of these positive regulators appears to be a distinguishing feature of the drought-tolerant species.

Next, we tried to identify TFs that probably regulate the elevated expression of these genes in the tolerant species. Three candidates, namely MYB73, cold shock domain protein 1 (CSDP1), and ERF4, bind multiple promoters in both the T0 subnetworks of *Aly* and *Esa* but are unexpectedly expressed at higher basal levels in *Ath*. All three have been characterized as negative regulators of abiotic-stress tolerance. Loss-of-function of *MYB73* enhances salt tolerance, whereas overexpression compromises survival under salt stress^51^. *CSDP1* overexpression delays seed germination under dehydration and salt stress, suggesting a repressive role in tolerance^52^. Similarly, overexpression of *ERF4* reduces ABA sensitivity and renders plants hypersensitive to NaCl, indicating that ERF4 dampens the salinity tolerance^53^. ERF4’s repressive role is further supported by our tobacco transactivation assay, in which it bound and downregulated the expression of the *Esa*-*MYC2* promoter (Fig. S1c). Taken together, these observations imply that the lower expression of *MYB73*, *CSDP1*, and *ERF4* in *Aly* and *Esa* may relieve repression of key stress-response genes, while their higher expression in *Ath* constrains such genes. Thus, low basal level expression of negative regulators of tolerance appears to be a hallmark of the drought-tolerant species.

Conversely, one TF in the T0 subnetworks, *Homeobox protein 6 (HB6)*, is expressed at higher basal levels in *Aly* and *Esa* than in *Ath.* HB6 binds three of the four shared high-basally expressed promoters: *MYC2, NADP-ME1*, and *SPL13A*, all of which act as positive regulators of tolerance. pampDAP assays showed that HB6 proteins from *Ath, Aly,* and *Esa* bind their respective *MYC2* promoters at similar positions (Fig. 5b), indicating a conserved interaction. In the tolerant species, the high basal level of *MYC2* (T0 log_2_FC > 4 relative to *Ath*) may be reinforced by (i) elevated basal expression levels of *HB6* and (ii) reduced expression levels of the negative regulators *ERF4, MYB73,* and *CSDP1.* Supporting this view, HB6 motifs are significantly enriched in the broader set of high basally expressed genes in *Aly* (P = 7.52 × 10^-06^, Fisher’s exact test) and *Esa* (P = 4.04 × 10^-02^, Fisher’s exact test) (Fig. 5d). HB6 is a transcriptional activator that is normally auto-induced by ABA, and overexpression in *Ath*, confers ABA insensitivity and increases water loss^54^. We propose that sustained HB6 expression in *Aly* and *Esa* provides a tunable ‘ABA desensitization’ mechanism, allowing tolerant species to temper an otherwise strong ABA sensitivity and thus balance growth with drought stress tolerance.

## Discussion

Our comparative analysis of drought-responsive gene regulatory networks in *Ath*, *Aly,* and *Esa* provides, to our knowledge, the most extensive cross-species resource of experimentally validated TF-promoter interactions under water-deficit stress to date. Nonetheless, several methodological constraints must be considered when interpreting our results.

We initially targeted 130 three-way ortholog gene promoters for Y1H. However, technical bottlenecks in PCR and cloning restricted the final dataset to 100 promoters (representing 36 ortholog sets), with complete three-species coverage for 29 sets. Although this corresponds to only 28% of the initially targeted candidates, the cloned orthologs captured 81% of the two-time point expression clusters (U/N/D based) present in the comparative drought transcriptome. Hence, despite this limited ortholog promoter coverage, our dataset remains sufficiently comprehensive to identify the transcriptional regulators of species-specific drought response.

Each promoter was confined to a window of -1.3 kb to +0.3 kb relative to the start codon; consequently, the effects of distal cis-elements and enhancers remain unexplored. However, genome-wide ATAC-seq across 13 angiosperms including *Ath* and *Esa* has shown that accessible chromatin is strongly concentrated at transcriptional start sites, with distal accessible regions accounting for only 6% of all accessible regions in *Ath*^55^. Thus, while distal cis elements may play more prominent roles in species with larger genomes, the proximal window employed here in *Ath, Aly,* and *Esa* with compact genomes should have captured most of the regulatory activity. Additionally, we applied stringent counter-selection to remove TFs that showed non-specific binding to the pMW2 plasmid backbone, which reduced potential false positives. Despite this filtering, the retained network contains TFs from 28 distinct families, encompassing all major drought-relevant regulators such as bZIP, DREB, ERF, HD-Zip, MYB, NAC, and WRKY. Therefore, we consider this diverse set of TFs adequately suited for dissecting core stress-response pathways.

We used *Ath* TFs as surrogates for their *Aly* and *Esa* orthologs, assuming conserved TF DNA-binding specificities within Brassicaceae. Although the literature largely supports this assumption, mutations in *Aly* and *Esa* TF coding sequences could have altered their DNA-binding specificities. As our analysis was limited to 1:1 orthologs among 1,334 conserved TFs, we excluded species-specific TFs that may contribute to lineage-specific aspects of drought adaptation. This orthology-focused approach is consistent with our previous work^5^ and other comparative GRN studies and ensures comparability between the species, enabling robust inference of regulatory trends. Although our strategy cannot capture the potential roles of *Aly-* and *Esa-*specific TFs, in the absence of comprehensive TF collections for these species, it remains the most interpretable and scalable framework currently available.

Despite these limitations, our network of 1,137 high-confidence TF-promoter interactions reveals trends and nodes that underlie phenotypic plasticity in drought response. Although detailed functional validation was beyond the scope of this study, multiple patterns emerge that warrant further investigation. First, in the drought-tolerant *Aly* and *Esa*, GRNs display markedly higher connectance and nestedness. While such topologies stabilize ecological networks, their role in strengthening molecular GRNs is unclear. However, analogous to the ecological networks, a denser and more redundant circuitry may itself promote drought tolerance by accelerating and amplifying signal propagation. If so, this elevated network density could account for some of the early or constitutive expression of stress protective genes in *Aly* and *Esa*. In this context, shifts in GRN connectance and nestedness may underlie the divergent stress response among closely related species.

Second, although *Aly* is phylogenetically closer to *Ath*, its drought-responsive GRN more closely mirrors that of *Esa*. This network-level similarity aligns with the comparable drought-survival rates of *Aly* and *Esa*, although their temporal expression dynamics under stress differ. Concurrently, similar patterns have been reported in distantly related drought-adapted plants, notably in desiccation-tolerant grasses, where recurrent environmental pressure drives GRNs toward a shared architecture regardless of lineage^56–58^. Collectively, these findings suggest that systems-level convergent evolution of GRNs shapes differential drought tolerance in both closely and distantly related species.

Third, we observed a genome-wide expansion of bZIP recognition motifs (G-boxes) in *Esa* relative to *Ath*, reflected by the increased fold enrichment of GBF3 and bZIP30 binding peaks on *Esa* promoters. We suspect that this expansion stems from transposable elements (TEs). *Esa’*s genome is nearly twice the size of *Ath*’s, and is composed of 52% of TEs (versus 13% in *Ath*), with Copia and Gypsy families predominating^46^. Many of these TEs are activated under environmental stress and carry stress-responsive TF binding sites^59,60^. Coupled with *Esa*’s lineage-specific expansion in ABA-biosynthetic pathway genes^46^, the additional G-boxes offer both more targets and binding sites for stress-activated TFs, representing a potential, albeit untested, driver of *Esa*’s improved drought tolerance. By contrast, extensive TE loss during *Ath*’s genome contraction may have pruned many such motifs from stress-gene promoters and contributed to its increased sensitivity to drought.

Basal expression profiles add yet another layer of contrast among the three species. *Aly* and *Esa* maintain low constitutive levels of the negative tolerance regulators *ERF4, MYB73*, and *CSDP1*, a state that likely heightens their sensitivity to ABA. In *Ath*, by contrast, higher basal expression of these TFs may hinder ABA-mediated activation, as reflected in its delayed T14 transcriptional response. Together with the G-box site expansion described above, this opposing regulation of negative tolerance factors offers a coherent explanation for the divergent drought-response plasticity observed with *Ath*, *Aly,* and *Esa*.

Finally, our network pinpoints edge-level species variations that may modulate drought outcomes. The GBF3-*CYP707A1* promoter interaction appears to contribute to susceptibility in *Ath*, whereas ABF-*mTERF10* promoter edges may enhance tolerance in *Esa*. Such effects are consistent with the broader principle that specific cis-regulatory element (CRE) changes can generate agronomically valuable traits during domestication^61^. For example, altered binding of OsSPL16 to a CRE in Grain width 7 produces more slender grains in rice^62^. Likewise, previous work showed that ABF binding increases transcript abundance; a single nucleotide change in the core ABF motif abolished binding and reduced expression of a stress-protective molecular chaperone^23^. These cases illustrate how gain or loss of individual edges, via CRE turnover, could help explain the contrasting drought tolerances seen across Brassicaceae species.

Beyond these trends, we validated ASIL2 as a novel regulator of drought response and identified additional promising candidates that warrant investigation. These include: *Hypothetical protein 1*, whose promoter showed the strongest regulatory similarity between *Aly* and *Esa*; bZIP30, which gained multiple *Esa*-specific interactions, and showed high pampDAP enrichment on *Esa* promoters; and *OFP7*, present in both tolerant species T0 subnetworks. Systematic evaluation of these targets will further refine network-guided approaches to engineering drought-resilient crops. To conclude, our integrative pipeline: combining transcriptome with systematic GRN mapping, provides a transferable framework for dissecting stress-adaptive mechanisms in closely related plant species. As full-length TF/ ORFeome collections expand in crops, this framework will become broadly applicable, accelerating the identification of cis- and trans-elements that underpin differential stress tolerance between closely related species. Taken together, our study presents both a resource and a methodology that can catalyse further research toward the rational engineering of drought-tolerant crops; an essential step toward securing yields in a changing climate.

---

**Figure S1a:**
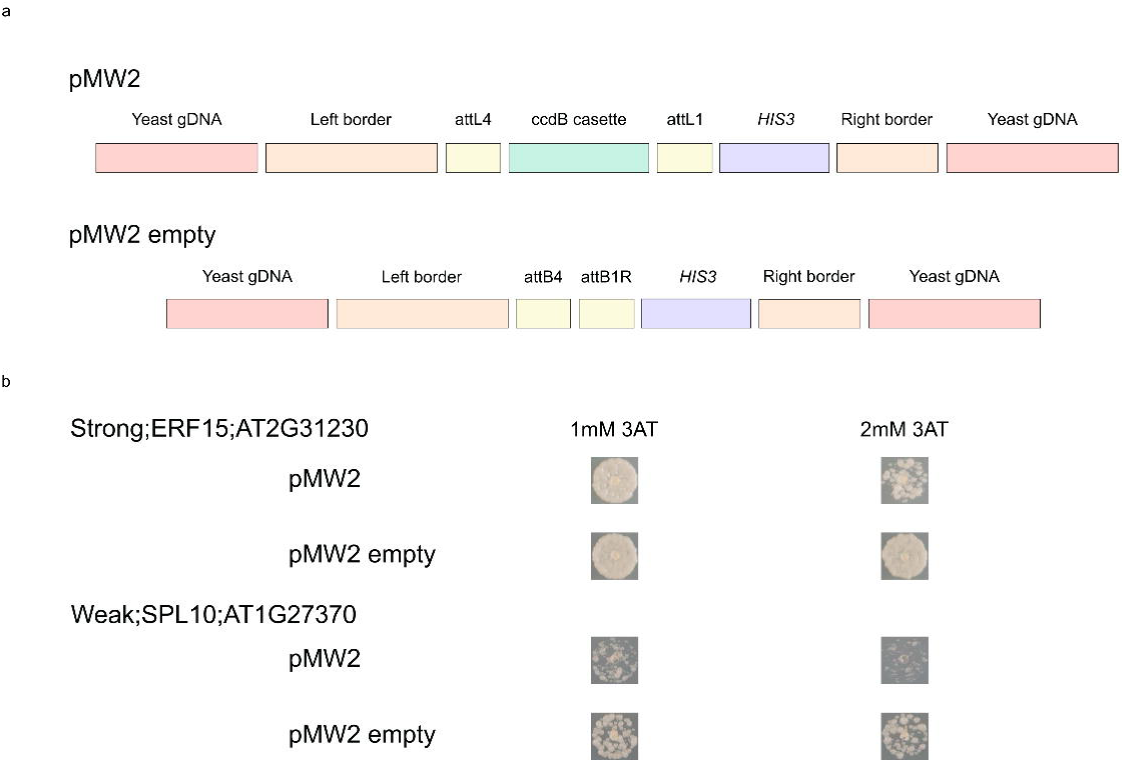
Identification of transcription factor autoactivators in Y1H. (a) Linearized map of plasmids used to identify TF autoactivators. (b) Y1H assay readout for weak and strong binding of TFs to pMW2 plasmid.

**Figure S1b:**
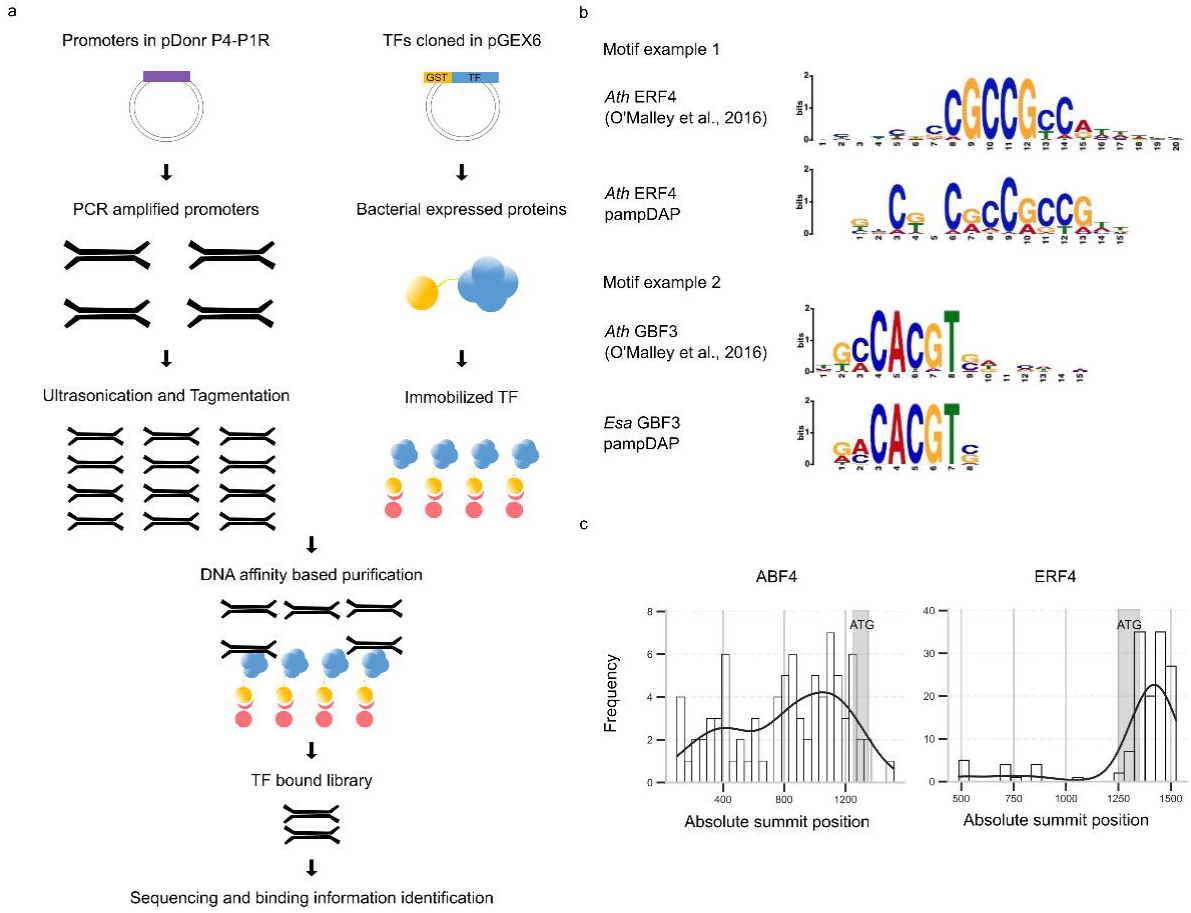
Promoter amplified DAP-seq workflow and motif analysis. (a) Overview of the pampDAP experimental workflow. (b) Comparisons of Literature-DAP and pampDAP motifs. (c) Distribution of pampDAP TF peak positions across the promoter length.

**Figure S1c:**
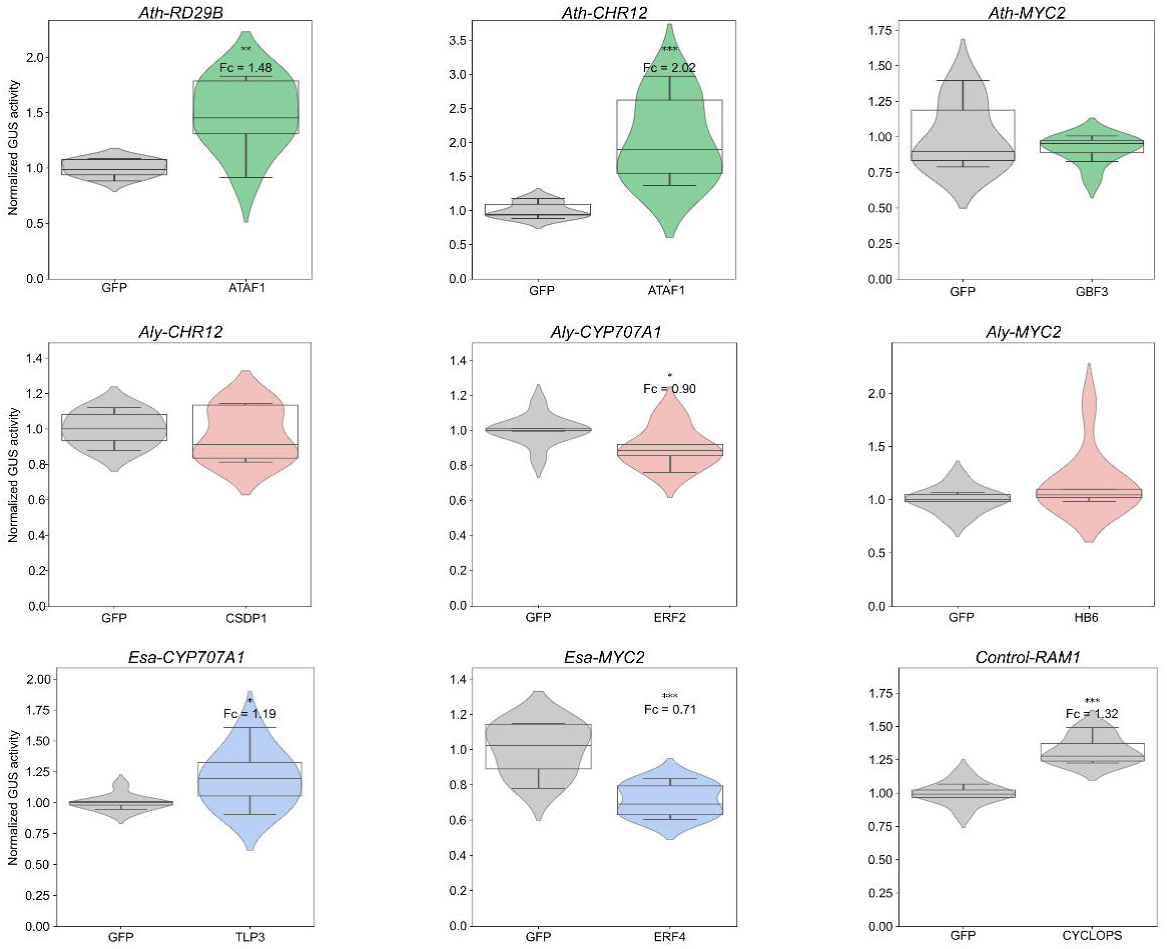
Validation of DGRN interactions in Tobacco transactivation assay. (***-p<0.05, one-sided Student’s t-test).

**Figure S1d:**
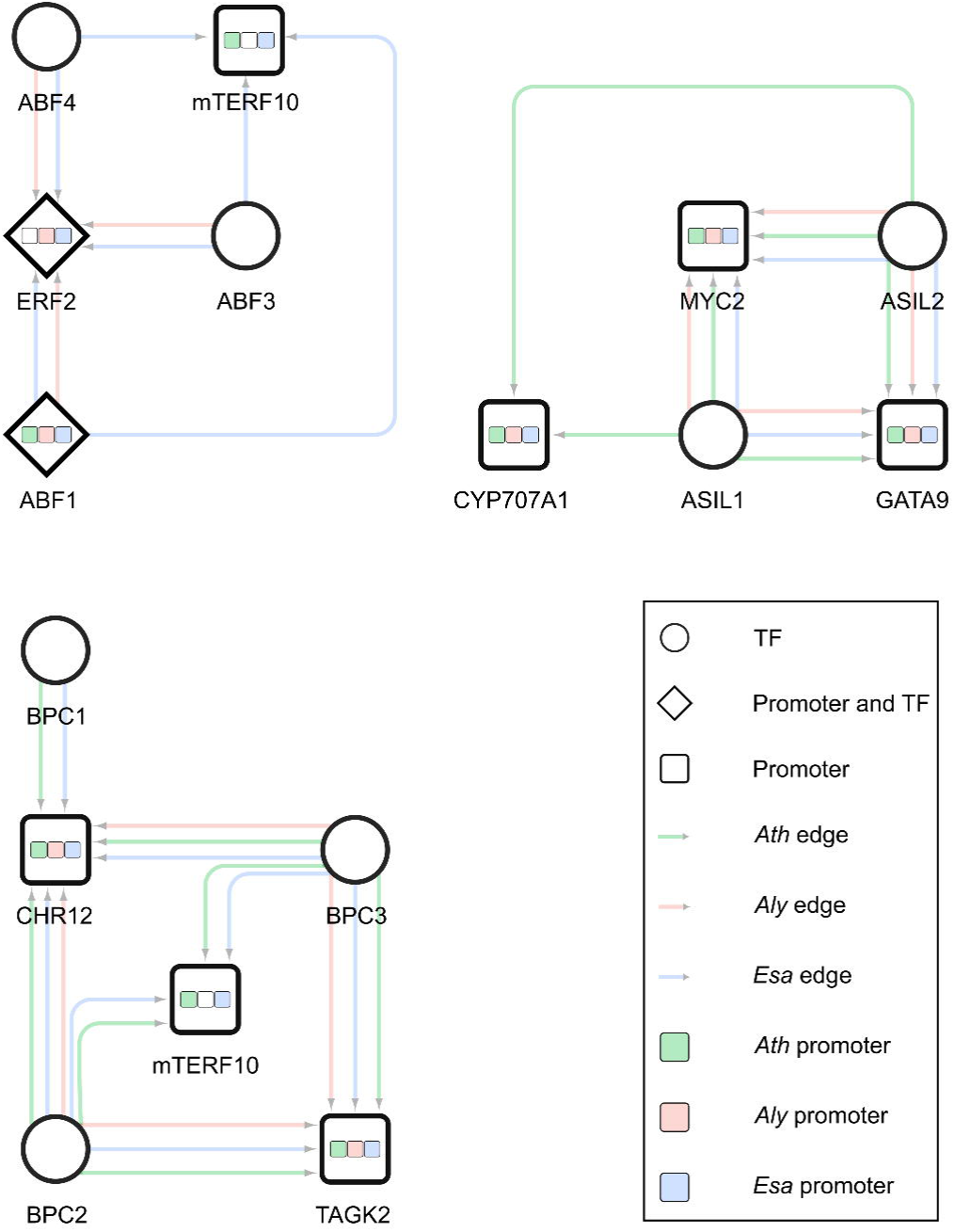
Redundant targeting of the same ortholog promoters by TF Family members.

## Author Contributions

R.P and P.F.B designed experiments. R.P, C.W.L, M.S, S.T.R, N.M.R, P.S, E.P, B.M, performed the experiments. R.P and P.F.B wrote the manuscript.

## Acknowledgments

ChatGPT was used to improve the text of the manuscript.

