## Supplementary material for "Comparative gene regulatory network mapping of Brassicaceae members with differential drought tolerance": Methods

### Experimental models

#### Bacteria

*E.coli* DH5α was used for cloning, and *E.coli* Rosetta strain was used for protein expression

#### Yeast

All yeast-one hybrid experiments were done using Y1HaS2 (*MATa ura3-52 ade2-101 ade5 lys2-801 leu2-3 his3-Δ1 trp1-901 tyr1-501 gal4D gal8D ade5::hisG*) and Yα1867 (*MATα SUC2 gal2 mal mel flo1 flo8-1 hap1 ho bio1 bio6 ura3-52 ade2-101 trp-901 his3-Δ200* ) strains

#### Plants

For experiment and cloning purposes, *Arabidopsis thaliana* (*Ath*) Col-0 ecotype, *Arabidopsis lyrata* (*Aly*) strain MN47, and *Eutrema salsugineum* (*Esa*) accession Shandong were used. Tobacco transactivation assays were done on *Nicotiana benthamiana* plants.

### Experimental methods

#### Promoter scoring and selection

We utilized the 15,883 orthologs established in our previous study[1] for expression divergence analysis. Each ortholog was assigned to one of three expression clusters at each time point-upregulated (U), not regulated (N), or downregulated (D)- based on fold-change (Fc) thresholds between successive time points: genes with Fc $\geq 1.5$ were classified as U, those with Fc $\leq 0.75$ as D and the remainder as N. Using this criterion across T5 and T11 transcriptome, we obtained categorical expression clusters (U/N/D) for all orthologs in *Ath*, *Aly*, and *Esa*.

To identify divergent expression patterns, we compared *Aly* and *Esa* clusters with *Ath* and introduced a metric called cluster distance to quantify expression shifts. A distance of 0 was assigned when cluster identity was conserved (e.g., U to U). Transitions between up/down-regulated (U/D) and non-responsive (N) clusters (e.g., U to N or D to N) were assigned a distance of 1. Direct opposing transitions (e.g., U to D or D to U) were assigned a distance of 2. For each ortholog, the maximum divergence score was 4, accounting for expression changes across two time points.

To identify species-specific and shared divergence, we constructed three scoring lists: (1) *Aly*-specific, (2) *Esa*-specific, and (3) *Aly-Esa* shared. For the *Aly-Esa* shared group, only genes showing identical U/N/D expression clusters in both *Aly* and *Esa* at both time points were retained to capture shared expression divergence.

Next, we annotated genes into three categories: transcription factors (TFs), signalling factors, and miscellaneous. TFs were annotated using established TF databases[2–4]. Genes encoding kinases[5], epigenetic regulators[6], phosphatases[7], ubiquitin ligases[8], fbox proteins[9] , co-activators[10] , and co-repressors[11] were grouped under regulators. All other genes that did not fall into these categories were labelled as miscellaneous.

To quantify expression magnitude divergence relative to *Ath*, we calculated the Manhattan distance between normalized expression values for both time points (T5 and T11), scaling the score between 0-4 to match the cluster distance scale. We then summed both metrics (Cluster distance + Manhattan distance) to compute a final divergence score for each gene.

For each of the three lists (*Aly*, *Esa*, *Aly-Esa*), we selected the top 50 most divergent genes within each functional category (TFs, signalling factors, miscellaneous). These were then merged to form three master lists. Each combined list was subjected to K-means clustering to identify distinct expression patterns, ensuring at least one representative gene was selected per cluster.

To prioritize regulatory components, we enforced a selection ratio of 50% TFs, 20% regulators, and 30% miscellaneous genes. Genes previously reported in our earlier study[1] and those involved in the ABA core pathway (biosynthesis, degradation, TFs, receptors, kinases, phosphatases, and epigenetic regulators) or associated with ABA/proline accumulation[12] [13] mutants were also included.

For the final set of 130 orthologs selected for promoter activity screening, primers were designed to amplify the -1300 + 300 region relative to the start codon (ATG) using the Primer-BLAST tool. For orthologs that failed this automated pipeline, we performed manual region walking to capture an equivalent region. This method was utilized for *OFP16* (AT2G32100), *FBS1* (AT1G61340), *MYB34* (AT5G60890), *MYC2* (AT1G32640), *PGIP1* (AT5G06860),

*AXR3* (AT1G04250), *DUF506* (AT2G20670), *CYP707A1* (AT4G19230), *BFT* (AT5G62040), *MYB112* (AT1G48000), *MAPKKK17* (AT2G32510), *PDF1.2C* (AT5G44430), *NAC19* (AT1G52890), *WRKY40* (AT1G80840), *AGL12* (AT1G71692), and *RBB1* (AT5G40450) promoters.

**Promoter cloning**

Promoter regions were PCR-amplified from genomic DNA of the respective species. In a second PCR, Gateway attB4 and attB1r recombination sites were added to them. The resulting products were recombined into the pDONR P4-P1r entry vector via a BP reaction (Invitrogen). Positive clones were confirmed by colony PCR and Sanger sequencing. Verified promoter entry clones recombined into the pMW2 destination vector using the Gateway LR reaction. Final destination plasmids were validated by PCR for correct promoter insertion.

**Yeast transformation and Y1H interaction screening**

pMW2 destination plasmids containing promoter sequences were linearized and integrated into the genome of the Y1HaS2 yeast strain following a previously published protocol[14]. To assess promoter autoactivation, transformed yeast bait strains were tested in a diploid format, more representative of screening conditions, by mating with the Yα1867 strain carrying a GFP reporter. For each promoter, 12 independent colonies were screened for autoactivation on Sc-His-Trp medium supplemented with 1,2, or 5mM 3-amino-1,2,4-triazole (3 AT). The colony exhibiting the lowest autoactivation was selected for further Y1H screening.

TF prey yeast strains were generated using a previously published TF collection[15]. For pooled screening, the entire collection (originally in 20 plates) was robotically compressed into two pool libraries, each comprising TFs from 10 plates. Pools were created by mixing 15µL from each source well into a single destination well, resulting in two 96-well TF pool plates.

Promoter bait strains were distributed into two 96-well plates and mated against each of the two TF pool plates. Mated diploids were selected and screened on SC-His-Trp medium supplemented with 1/2/5 mM 3AT, depending on the promoter's autoactivation profile. Interactions were scored using a custom machine learning-based image analysis pipeline[16]. A pooled spot was considered positive if colony growth exceeded the median growth on the corresponding plate. Colonies from positive spots were picked and sequenced using next-generation sequencing (NGS) to identify interacting TFs[17]. Identified TFs were then rearrayed and retested individually to confirm interactions from orthologous promoters.

For focused Y1H screening, 188 TFs were selected based on Gene Ontology (GO) terms relevant to drought responses, such as “response to abiotic stimulus,” “response to radiation,” ”response to osmotic stress,” “response to water deprivation,” and “ response to ABA.” Promoter strains were mated once, with the TF prey strains in two 96-well plates. Interactions identified via the machine learning model were subsequently validated in triplicate using rearrayed prey strains.

To identify TFs that may interfere nonspecifically with the assay setup, both the empty pMW2 vector and the full pMW2 backbone (with ccdB region) were screened in 2 and 5 mM 3AT. TFs that interacted strongly with both plasmids were removed from the dataset.

**pampDAP assay**

The TFs from Gateway--compatible entry plasmids, were transferred into a Gateway-adapted pGEX-6 plasmid containing an N-terminal GST tag using the LR reaction. All TF inserts were verified by colony PCR, followed by plasmid minipreps and sequence confirmation.

The resulting expression constructs were transformed into *E.coli* Rosetta (DE3) cells for protein expression. Cultures were grown in LB supplemented with appropriate antibiotics and induced with 1mM IPTG at mid-log phase ($OD_{600}$ = 0.6) for 3hours at 37°C. GFP was used as an expression and purification control.

Cells were harvested by centrifugation, lysed, and recombinant proteins were purified using the MagneGST™ protein purification system (Promega), following the manufacturer’s instructions. Protein purity and integrity were assessed via SDS-PAGE, and protein concentrations were quantified using the Pierce™ BCA protein assay kit (Thermo Scientific). Size-based purification was performed using Amicon Ultra 0.5 ml centrifugal filter devices (Millipore).

Promoters cloned into pDONR-P1r were PCR-amplified using custom-designed primers. PCR products were purified with magnetic beads and quantified using the Quant-iT™ PicoGreen dsDNA assay kit (Invitrogen). Equal volumes (5 µl) from each purified PCR product were pooled to generate a promoter DNA library.

The pooled promoter library was adjusted to a final concentration of 100ng/µl in elution buffer (5 mM Tris-HCl, pH 8.5) and was fragmented using a Covaris E220 focused ultrasonicator to obtain DNA fragments ranging from 250-500 bp. Fragmented DNA was then subjected to tagmentation using Illumina-Tn5 transposase, followed by PCR amplification using indexed Illumina adapters.

Final libraries were size-selected (250-500bp), purified, and quality-checked before sequencing. DNA-affinity purification, peak calling, and motif analysis were conducted as described previously[18]. The entire experiment was performed in biological duplicates, and unique peaks identified in both replicates were retained for downstream analyses.

**Tobacco transactivation assay**

To assess the effect of TF interactions on promoter activity, we built a custom transient transactivation assay in tobacco. A modified version of the pGWB3 binary vector was constructed to serve as the promoter-reporter backbone[19]. The Gateway cassette in pGWB3 was excised using HindIII and Sma1, and replaced with a synthetic “dummy” sequence that introduces a proximal Sma1 site upstream of the GUS coding sequence, generating the reporter vector P1H.

Promoter sequences were PCR-amplified from the Y1H destination plasmid (pMW2) using 5’-phosphorylated primers containing a CaMV 35s minimal promoter sequence. The resulting promoter fragments were ligated into the Sma1-digested, dephosphorylated P1H plasmid using blunt-end ligation. The correct insertion orientation of promoters in bacterial colonies was verified via colony PCR.

TF open reading frames were cloned under the control of the CaMV 35s promoter into the pGWB2 vector using the LR Clonase reaction. All final constructs were mini-prepped from *E.coli*, transformed into *Agrobacterium tumefaciens* strain GV3101, and confirmed via colony PCR.

For leaf infiltration, *Agrobacterium* cultures harboring the constructs were grown for 48 hours, washed, and resuspended to a final $OD_{600}$ of 1.0 in infiltration buffer (10 mM $MgCl_2$, 10 mM MES pH 5.6, 150 μM Acetosyringone), followed by a 2-hour incubation at room temperature.

Fully expanded leaves of 5-week-old tobacco plants were used for infiltration. Equal volumes of appropriate cultures were mixed to prepare infiltration combinations: Promoter::GUS + TF/GFP (negative control) + P19 (suppressor of silenceing). For each promoter-TF pair, the corresponding promoter-GFP control was always co-infiltrated within the same leaf. Each combination was tested in triplicate (technical replicates) within a leaf and repeated across multiple plants (biological replicates).

Two days post-infiltration, leaf discs were collected using a leaf puncher, flash frozen in liquid nitrogen, and stored at -80°C with glass beads. Samples were homogenized using a bead mill, and lysates were prepared by centrifuging at 4,500 rpm. 200μl of extraction buffer (50mM sodium phosphate pH 7.0, 10 mM EDTA pH 8.0, 0.1% SDS, 0.1% Triton X-100, 10 mM β-mercaptoethanol) was added, followed by a final centrifugation to remove debris.

10 μl of the clarified lysate was used for GUS fluorometric assay in 100 μl incubation buffer (50 mM sodium phosphate pH 7.0, 10mM EDTA pH 8.0, 0.1% Triton X-100, and 1 mM MUG) and incubated for 4 hours at 37°C. Reactions were stopped with 20 μl of 0.2M sodium carbonate, and fluorescence was measured.

Protein concentration was determined using the BCA assay (Thermo Fisher). GUS activity was normalized to total protein. To compare TF-dependent activity, GUS activity in the Promoter + TF condition was normalized to its paired Promoter + GFP negative control from the same leaf, and comparisons were then made across different biological replicates.

**Plant phenotype measurements**

T-DNA insertion mutant seeds were obtained from NASC, screened for homozygosity, and propagated for experimental use. Seeds were surface sterilized using bleach followed by 75% ethanol, then stratified and germinated on half-strength MS medium. After initial growth, seedlings were transferred to plates containing either 0mM or 25mM mannitol. Photos of the seedlings were captured 7 days after the transfer.

Projected Rosette Area (PRA) was calculated using ImageJ software. PRA values were normalized to the wild type (Col-0) growth at 0 mM Mannitol within the same repeat. Growth reduction in the mutant was assessed relative to the wild-type response under osmotic stress.

### Network analysis and figure generation

Gene regulatory networks were visualized using Cytoscape V3.10.3[20]. Figures were generated using Excel, R (ggplot2), and Python (matplotlib and seaborn) packages with support from Julius AI (https://julius.ai/).

For network topology analysis, the Bipartite package[21] in R was used to calculate metrics such as connectance and nestedness. Nestedness was calculated using NODF (Nestedness as a measure of Overlap and Decreasing fill) score. Coregulation scores between transcription factor-target gene pairs were calculated as the coefficient of determination[22] between expression profiles of interacting and non-interacting pairs.
